# Bimodal Regulation of the PRC2 Complex by USP7 Underlies Melanomagenesis

**DOI:** 10.1101/641977

**Authors:** Dongxue Su, Wenjuan Wang, Yongqiang Hou, Liyong Wang, Yue Wang, Chao Yang, Beibei Liu, Xing Chen, Xiaodi Wu, Jiajing Wu, Dong Yan, Shuqi Wei, Lu Han, Shumeng Liu, Lei Shi, Lin Shan

## Abstract

Although overexpression of EZH2, a catalytic subunit of the polycomb repressive complex 2 (PRC2), is an eminent feature of various cancers, the regulation of its abundance and function remains insufficiently understood. We report here that the PRC2 complex is physically associated with ubiquitin-specific protease USP7 in melanoma cells where USP7 acts to deubiquitinate and stabilize EZH2. Interestingly, we found that USP7-catalyzed H2BK120 deubiquitination is a prerequisite for chromatin loading of PRC2 thus H3K27 trimethylation. Genome-wide analysis of the transcriptional targets of the USP7/PRC2 complex identified a cohort of genes including *FOXO1* that are involved in cell growth and proliferation. We demonstrated that the USP7/PRC2 complex drives melanoma cell proliferation and tumorigenesis *in vitro* and *in vivo.* We showed that the expression of both USP7 and EZH2 elevates during melanoma progression, corresponding to a diminished FOXO1 expression, and the level of the expression of USP7 and EZH2 strongly correlates with histological grades and prognosis of melanoma patients. These results reveal a dual role for USP7 in the regulation of the abundance and function of EZH2, supporting the pursuit of USP7 as a therapeutic target for melanoma.

## INTRODUCTION

Polycomb repressive complex 2 (PRC2), which includes EED, SUZ12, RbAp46, RbAp48, and the enzymatically active core subunit enhancer of zeste homolog 2 (EZH2), is required to maintain the transcriptional repression of key developmental regulators during normal embryonic development (1). EZH2 methylates H3K27, which facilitates chromatin compaction and gene silencing (2). Several lines of evidence have implicated EZH2 in the development and progression of a variety of cancers, and elevated expression of EZH2 often correlates with a poor prognosis (3, 4). An early indication of a role for EZH2 in prostate cancer was based on the observation that EZH2 overexpression is associated with worse progression of prostate cancer (5). Subsequently, similar findings emerged in other human cancers such as breast cancer (6), endometrial cancer (7), and melanoma (8). Thus, understanding the molecular basis of how dysregulation of EZH2 contributes to tumorigenesis is of great significance in cancer treatment and prevention.

Melanoma is the most aggressive form of skin cancer, and the incidence is increasing annually (9). Unlike localized melanoma, which can be surgically resected, metastatic melanoma with extranodal involvement is typically treated with systemic therapies, such as targeted therapy. Vemurafenib, a BRAF inhibitor, has been shown to improve outcomes in the majority of melanoma patients harboring the highly prevalent BRAF V600E mutation (10). However, most patients treated with vemurafenib show disease progression within 6–8 months due to invariable drug resistance (11). Thus, identification of novel theraputic targets is urgently needed.

BRAF, located on chromosome 7q34, is adjacent to the EZH2 gene on chromosome 7q36.1 (12, 13). Indeed, abnormalities in these two genes often coexist in various types of cancers (14). It is well-documented that EZH2 is highly expressed in human melanoma, which is linked to poor patient survival (14). Mutations affecting EZH2 activity are predominantly found in melanoma (15) and its gain-of-function mutations often occur concurrently with the BRAF V600E mutation (14). This suggests a close relationship between EZH2 function and BRAF mutation, thus highlighting EZH2 as a promising combination therapeutic target with vemurafenib for melanoma therapy.

Ubiquitination levels are balanced by ubiquitinating enzymes, including E1, E2, and E3 (16), and deubiquitinating enzymes (DUBs) (17). Among the known DUBs, ubiquitin-specific protease 7 (USP7), also known as herpes virus-associated ubiquitin-specific protease (HAUSP) (18) since it was originally identified as a herpes simplex virus type 1 Vmw110-interacting protein (19), is associated with multiple cellular processes such as DNA repair (20), innate immune responses (21), viral replication and infection (22), and mitosis progression (23). In addition, we recently reported that USP7 promotes breast carcinogenesis by stabilizing PHF8 and upregulating several key cell cycle regulators such as cyclin A2 (24). USP7 was also found to stabilize mediator of DNA damage checkpoint protein 1 (MDC1), which promoted cervical cancer cell survival and conferred cellular resistance to genotoxic insults (25). Although elevated *USP7* expression was recently reported as a distinct gene signature of malignant melanoma (26), its specific contribution to melanoma progression is largely unexplored.

Ubiquitination also occurs on histones, especially H2A and H2B (27). Mono-ubiquitination of H2A and H2B impacts a wide range of chromatin-associated events such as transcription, chromatin organization, and DNA replication (28). Dysregulation of H2A or H2B ubiquitination has been associated with several pathological processes, particularly cancer (29, 30). Analogously, the state of histone ubiquitination is balanced by the activities of the specific ubiquitin ligases and deubiquitinases. In *Drosophila* testes, it was demonstrated that USP7 formed a protein complex with guanosine 5’-monophosphate synthetase to catalyze the removal of H2BK120ub1 (31). It seems that H2BK120ub1 and histone H2A K119 monoubiquitylation (H2AK119ub1) are mutually exclusive, since these two histone marks are associated with transcriptional activation and repression, respectively. As H2AK119ub1 and trimethylation of histone H3 at K27 (H3K27me3) are considered to be functionally coordinated in transcriptional repression (32), it will be interesting to investigate the detailed molecular mechanism underlying the coordination between H2BK120ub1 deubiquitination and H3K27me3.

FOXO family factors have been shown to have a key role in a variety of biological processes, including cell cycle (33), apoptosis (34), oxidative and stress response (35, 36). Among them, FOXO1 involved in a wide range of organismal functions, especially function as a tumor suppressor (37–39). It has been reported that dis-regulation of FOXO1 is associated with multiple cancers, such as prostate cancer (40), breast cancer (41), and endometrial cancer (38). However, the relationships between clinical significance and FOXO1 expression as well as the underlying mechanisms of FOXO1 expression in melanoma have not been established.

In this study, we revealed that deubiquitinase USP7 is physically and functionally associated with the PRC2 complex. Specifically, we demonstrated that, on one hand, USP7 deubiquitinates and stabilizes EZH2, on the other hand, USP7-promoted H2BK120ub1 removal dictates PRC2 recruitment thus methylation of H3K27. In this manner, USP7 coordinates with the PRC2 complex to control the expression of tumor suppressors, including FOXO1 to promote melanoma proliferation *in vitro* and *in vivo*.

## RESULTS

### USP7 Is Physically Associated with PRC2 Complex

In an effort to better understand the biological function of EZH2 and to investigate its role in the development of cancer, we generated a human malignant melanoma A375 cell line, which can stably express FLAG-EZH2. Whole-cell extracts from these cells were subjected to affinity purification using an anti-FLAG affinity gel. Mass spectrometric (MS) analysis indicated that EZH2 was associated with SUZ12 and EED, subunits of the PRC2 complex. Additional proteins, including OGT and SPTBN1, were also detected in the EZH2-containing complex. Interestingly, USP7 was also identified in the EZH2-containing protein complex with high peptides coverage (Figure 1A and Supplementary Table 1). To further confirm the physical interaction between EZH2 and USP7, we generated an A375 cell line with doxycycline (Dox)-inducible expression of stably integrated FLAG-USP7. Similarly, the MS analysis of the whole-cell extracts from these cells with or without Dox-inducible expression of FLAG-USP7 showed that the deubiquitinase USP7 was indeed co-purified with PRC2 complex (Figure 1A and Supplementary Table 1).

**Figure 1.**
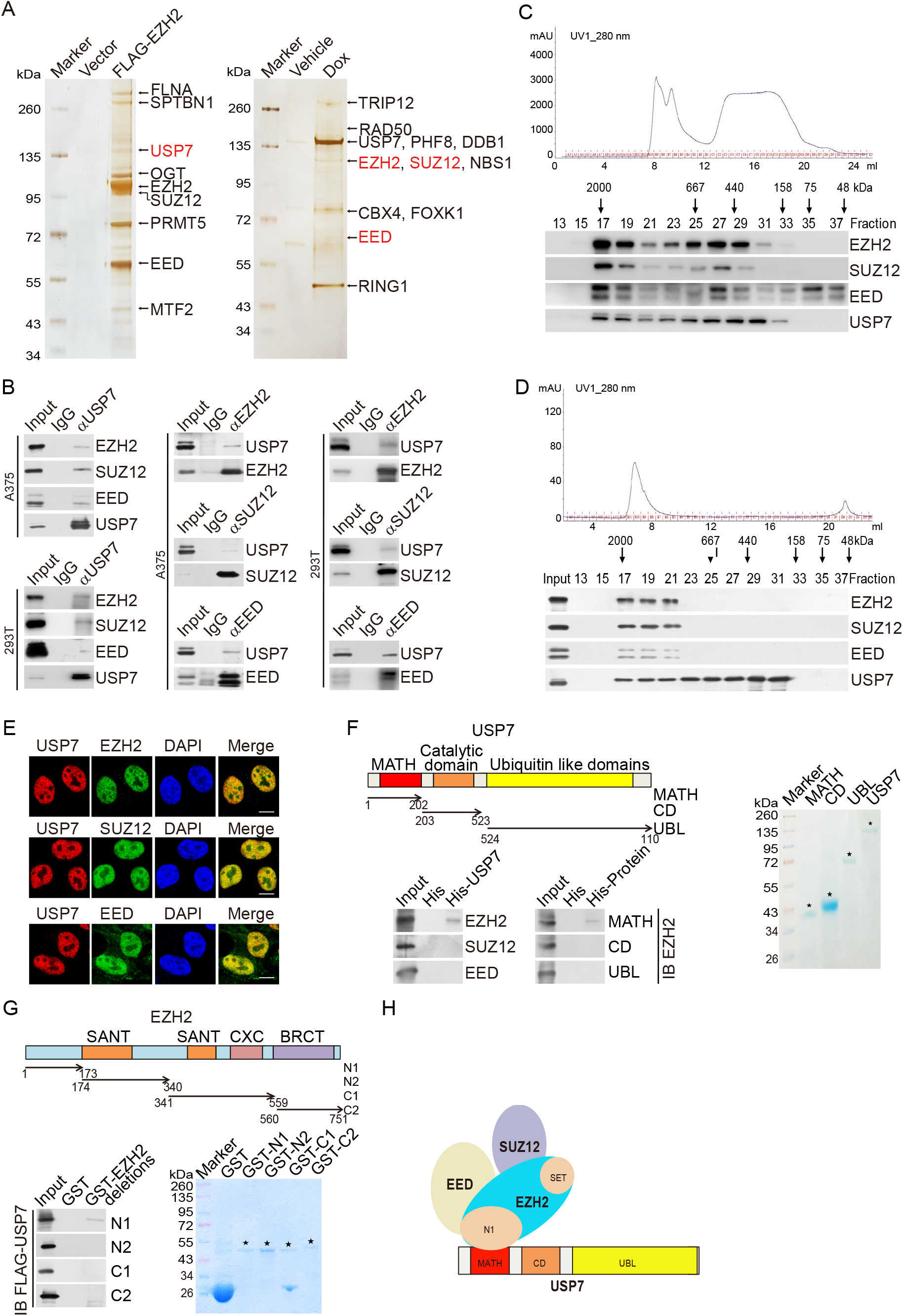
USP7 is physically associated with PRC2 complex. (A) Immunopurification and mass spectrometry analysis of EZH2- and USP7-containing protein complex. Cellular extracts from FLAG-EZH2 or FLAG-USP7-stably expressing A375 cells were immunopurified with anti-FLAG-affinity beads and eluted with FLAG peptide. The eluates were resolved on SDS/PAGE and silver-stained followed by mass spectrometry analysis. (B) Co-IP analysis of the association between USP7 and PRC2 complex. Whole cell lysates from A375 cells and HEK293T cells were immunoprecipitated (IP) then immunoblotted (IB) with antibodies against the indicated proteins. (C) Fast protein liquid chromatography analysis of the native protein complex. Chromatographic elution profiles (upper panel) and western blotting analysis (lower panel) of the chromatographic fractions with antibodies against the indicated proteins are shown. Equal volumes from each fraction were analyzed and the elution positions of calibration proteins with known molecular masses (kDa) are indicated. (D) Experiments analogous to (C) were performed with the USP7-containing protein complex purified from FLAG-USP7-expressing A375 cells. (E) Confocal microscopy analysis of USP7 and PRC2 complex subcellular localization. A375 cells were fixed and immunostained with antibodies against the indicated proteins. Scale bar, 10 μm. (F) His-pull down assays with full-length or deletion mutants of USP7 purified from Sf9 cells and *in vitro*-transcribed/translated proteins as indicated. (G) GST pull-down assays with bacterially expressed GST-fused deletion mutants of EZH2 and *in vitro-*transcribed/translated FLAG-USP7. (H) Illustration of the molecular interfaces required for the association of USP7 with the PRC2 complex.

To confirm the *in vivo* interaction of USP7 with the PRC2 complex, total proteins from A375 and 293T cells were extracted, and co-immunoprecipitation (co-IP) was performed with antibodies detecting the endogenous proteins. Immunoprecipitation (IP) with antibodies against EZH2, SUZ12, and EED or USP7 followed by immunoblotting (IB) with USP7 or the PRC2 complex proteins, respectively, in A375 and 293T cells confirmed that endogenous PRC2 complex interacts with USP7 *in vivo* (Figure 1B).

To further support the physical interaction of USP7 with PRC2 complex, fast protein liquid chromatography (FPLC) with A375 nuclear extracts using a Superpose 6 column and a high salt extraction and size-exclusion approach further supported the physical interaction of USP7 with PRC2 complex. Native USP7 in A375 cells was eluted with an apparent molecular mass much greater than that of the monomeric protein, and the chromatographic profile of USP7 largely overlapped with that of EZH2, SUZ12, and EED (Figure 1C). Furthermore, the majority of the purified FLAG-USP7 existed in a multiprotein complex, which peaked in fractions 17-19 with PRC2 complex (Figure 1D). Consistently, immunofluorescent (IF) staining with antibodies against endogenous EZH2, SUZ12, EED and USP7 showed that PRC2 complex and USP7 were colocalized in the nucleus of A375 cells (Figure 1E).

To substantiate the physical interaction of USP7 with PRC2 complex *in vivo* and to understand the molecular details involved in this interaction, pull-down experiments with full-length USP7 purified from Sf9 cells and recombinant EZH2, SUZ12 or EED demonstrated that USP7 directly interacts with EZH2, but not with the other proteins tested (Figure 1F). USP7 contains three distinct structural domains: the C-terminal MATH domain, the catalytic domain (CD), and the N-terminal ubiquitin-like domain (UBL) (24). Incubation of recombinant EZH2 with truncation mutants of USP7 purified from Sf9 cells showed that MATH domain-containing truncation mutants of USP7 could directly interact with EZH2 (Figure 1F). Moreover, glutathione S-transferase (GST) pull-down with GST-fused EZH2 N1 domain (aa 1-173), EZH2 N2 domain (aa 174-340), EZH2 C1 domain (aa 341-559), EZH2 C2 domain (aa 560-751), and *in vitro*-transcribed/translated USP7 revealed that USP7 directly interacts with the N1 domain of EZH2 (Figure 1G). Together, these results not only provide further support of the specific interaction between USP7 and PRC2 complex but also delineated the molecular details involved in the formation of the USP7/PRC2 complex, as schematically summarized in Figure 1H.

### USP7 Is Functionally Linked to the Stability and Deubiquitination of EZH2

To investigate the functional significance of the physical interaction and spatial co-localization between USP7 and EZH2, we examined the effect of USP7 on the expression of EZH2. Western blotting analysis of cellular lysates from A375 cells transfected with three independent sets of siRNAs targeting different regions of *USP7* revealed that the level of EZH2 was significantly reduced upon USP7 depletion (Figure 2A), although quantitative reverse transcription PCR (qRT-PCR) indicated that USP7 knockdown did not result in alterations in *EZH2* mRNA levels in A375 cells (Figure 2A). Meanwhile, the effect was blocked by treatment of a proteasome-specific inhibitor, MG132 (Figure 2B). This suggests that the reduction in EZH2 protein level associated with USP7 depletion involves a proteasome-mediated protein degradation mechanism rather than transcription-level regulation. These observations indicated that the stability of EZH2 is regulated by USP7 and that EZH2 is a substrate of USP7.

**Figure 2.**
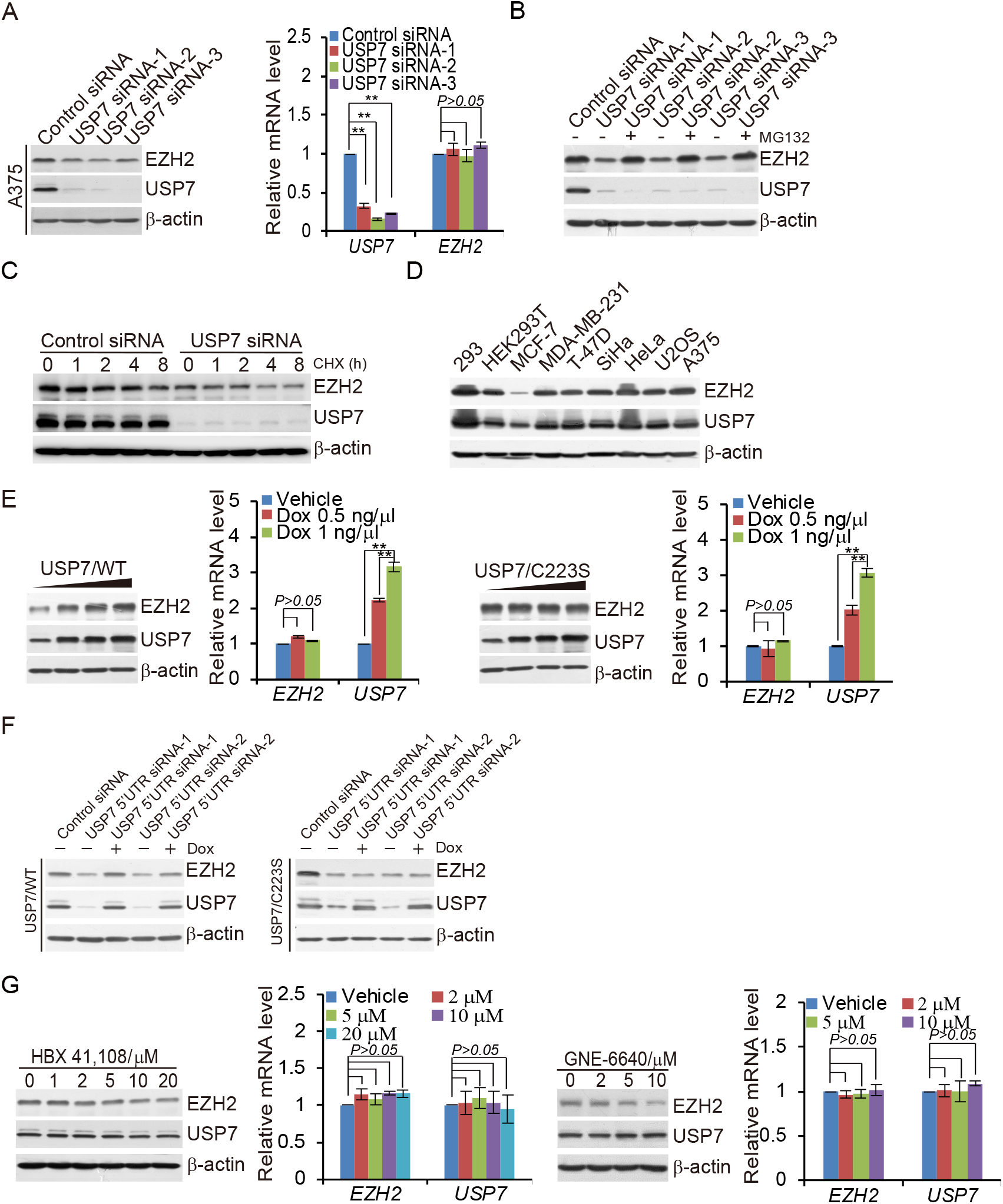
USP7 promotes EZH2 stabilization. (A) A375 cells transfected with control siRNA or different sets of *USP7* siRNAs were collected and analyzed by western blotting and quantitative reverse transcription (qRT)-PCR. Each bar represents the mean ± SD from biological triplicate experiments. \*\**P* < 0.01, by two-way ANOVA. (B) A375 cells transfected with control siRNA or *USP7* siRNAs were treated with DMSO or proteasome inhibitor MG132 (10 μM). Cellular extracts were prepared and analyzed by western blotting. (C) A375 cells were transfected with control siRNA or *USP7* siRNA followed by treatment with cycloheximide (CHX, 50 μg/ml), and harvested at the indicated time points followed by western blotting analysis. (D) Western blotting analysis of the expression of USP7 and EZH2 in multiple cell lines. (E) A375 cells with Dox-inducible expression of FLAG-USP7/WT or FLAG-USP7/C223S cultured in the absence or presence of increasing amounts of Dox. Cellular extracts and total RNA were collected for western blotting and qRT-PCR analysis, respectively. Each bar represents the mean ± SD from biological triplicate experiments. \*\**P* < 0.01, two-way ANOVA. (F) A375 cells with Dox-inducible expression of FLAG-USP7/WT or FLAG-USP7/C223S transfected with control siRNA or different sets of *USP7* 5′UTR siRNAs in the absence or presence of Dox. Cellular extracts were collected and analyzed by western blotting. (G) A375 cells cultured in the absence or presence of increasing amounts of HBX 41,108 for two hours or with GNE-6640 for 24 hours as indicated. Cellular extracts and total RNAs were collected for western blotting and qRT-PCR analysis, respectively. Each bar represents the mean ± SD from biological triplicate experiments. *P* values were calculated by one-way ANOVA.

To further support this deduction, the potential of USP7 to modulate the steady-state level of EZH2 protein was assessed by cycloheximide (CHX) chase assays. After incubation with CHX, western blotting analysis revealed that USP7 depletion was clearly associated with a decreased half-life of EZH2 (Figure 2C). In addition, the expression levels of USP7 and EZH2 were correlated in multiple cell lines (Figure 2D).

To further support the relationship between USP7 and EZH2, we next investigated whether USP7-promoted EZH2 stabilization is dependent on the enzymatic activity of USP7. To this end, we generated two stable A375 cell lines, wild type USP7 (USP7/WT) and a catalytically inactive mutant of USP7 (USP7/C223S), both with Dox-inducible expression. Western blotting analysis showed a significant increase in the protein level of EZH2 in a Dox dose-dependent manner in cells expressing USP7/WT, but not USP7/C223S, while there was no change in the mRNA level of *EZH2* in either cell line (Figure 2E). Moreover, the downregulation of EZH2 in USP7-deficient cells could be reverted by forced expression USP7/WT, but not USP7/C223S (Figure 2F). Treatment of A375 cells with HBX41,108, a deubiquitinase inhibitor that inhibits the catalytic activity of USP7, or GNE-6640, a recently developed USP7 inhibitor, reduced the protein level of EZH2 but did not affect the *EZH2* mRNA level (Figure 2G).

To further support the notion that USP7 is the deubiquitinase of EZH2, A375 cells stably expressing FLAG-EZH2 were co-transfected with USP7 siRNA and HA-tagged wild type ubiquitin (Ub/WT). IP of the cellular lysates with anti-FLAG followed by IB with anti-HA showed that knockdown of USP7 increased the level of ubiquitinated EZH2 species (Figure 3A). Moreover, enzymatic inhibition of USP7 with HBX 41,108 or GEN-6640 resulted in a marked increase in the level of ubiquitinated EZH2 species (Figure 3B). In addition, A375 cells stably expressing FLAG-EZH2 were co-transfected with different amounts of USP7 and HA-Ub/WT or an HA-tagged ubiquitin mutant (Ub/MT) with all lysine resides replaced by arginine. IP of cellular lysates with anti-FLAG followed by IB with anti-HA showed that increased USP7 expression was associated with decreased levels of ubiquitinated EZH2 species (Figure 3C). However, there were no changes in the levels of ubiquitinated EZH2 species with increasing amounts of USP7/C223S expression (Figure 3D). To further support this proposition, A375 cells stably expressing FLAG-EZH2 were co-transfected with *USP7* siRNAs and HA-tagged Ub/K48-only, a ubiquitin mutant with all lysine resides replaced by arginine except for residue 48 that signals targets proteins for degradation, or with HA-tagged Ub/K63-only, a ubiquitin mutant with all lysine resides replaced by arginine except for lysine 63, which is responsible for the proteasome-independent signals. USP7 was capable of deubiquitinating K48-linked polyubiquitin chains but not K63-linked polyubiquitin chains (Figure 3E). Together, these results indicate that USP7 could regulate the stability of EZH2 through its deubiquitinase activity.

**Figure 3.**
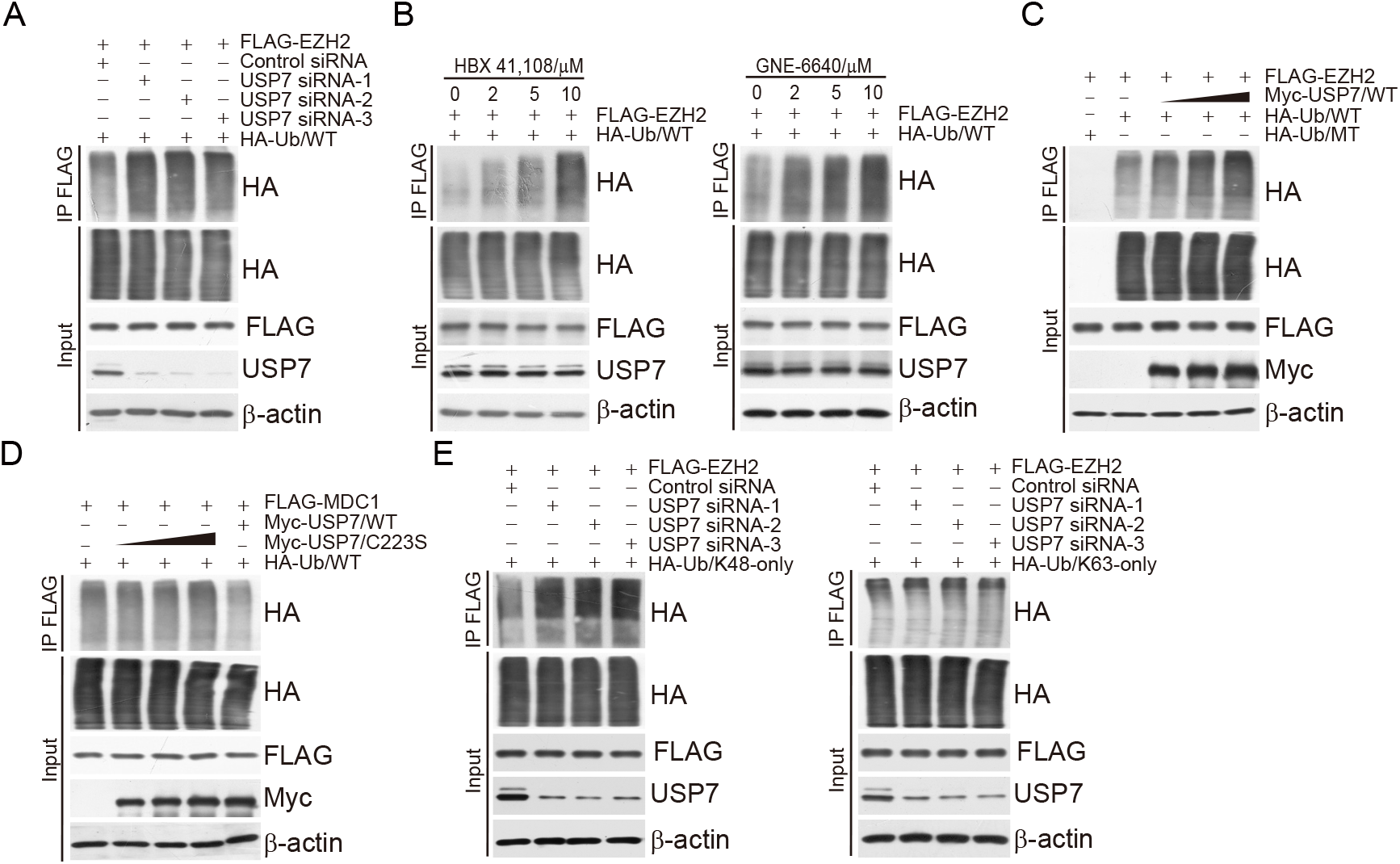
USP7 deubiquitinates EZH2. (A) A375 cells stably expressing FLAG-EZH2 co-transfected with control siRNA or *USP7* siRNAs and HA-Ub/WT. Cellular extracts were immunoprecipitated with anti-FLAG followed by immunoblotting with anti-HA. (B) A375 cells stably expressing FLAG-EZH2 transfected with HA-Ub/WT and cultured in the presence or absence of HBX 41,108 and GNE-6640. Cellular extracts immunoprecipitated with anti-FLAG followed by immunoblotting with anti-HA. (C) A375 cells stably expressing FLAG-EZH2 were co-transfected with HA-Ub/WT or HA-Ub/MT and different amounts of Myc-USP7. Cellular extracts were immunoprecipitated with anti-FLAG followed by immunoblotting with anti-HA. (D) A375 cells stably expressing FLAG-EZH2 co-transfected with HA-Ub/WT and Myc-USP7/WT or different amounts of Myc-USP7/C223S as indicated. Cellular extracts were immunoprecipitated with anti-FLAG followed by immunoblotting with anti-HA. (E) A375 cells stably expressing FLAG-EZH2 co-transfected with different amounts of Myc-USP7/WT and HA-Ub/K48-only or HA-Ub/K63-only followed by immunoprecipitation analysis.

### USP7-catalyzed H2BK120 Monoubiquitination Removal Is a Prerequisite for Chromatin Occupancy of PRC2 thus H3K27 Trimethylation

As PRC2 complex are well-known epigenetic regulators of transcription, and USP7 is believed to be a deubiquitinase for H2BK120ub1, we wondered whether USP7 plays a role in modulating the level of H3K27me3. To this end, A375 cells were transfected with USP7 siRNA. Forty-eight hours posttransfection, cells were fixed and stained with antibodies recognizing USP7, EZH2, H3K27me3, and H2BK120ub1, immunofluorescence staining followed by confocal microscopy analysis revealed that endogenous USP7 appeared to be colocalized with EHZ2; meanwhile, the signals representing EZH2 and H3K27me3 were significantly decreased in USP7-deficient A375 cells, while H2BK120ub1 signals were slightly elevated (Figure 4A). Next, to understand the functional interplays between USP7 and EZH2 in transcriptional repression in A375 cells, USP7 siRNA or EZH2 siRNA with FLAG-Vector or FLAG-EZH2 were co-transfected in the melanoma A375 cells. Western blotting showed that knockdown of USP7 in A375 cells resulted in a decreased level of H3K27me3, even when EZH2 was overexpressed (Figure 4B). Similar results were obtained in HaCat cells, a spontaneously transformed aneuploid immortal keratinocyte cell line from adult human skin, and human melanocyte PIG1 cells (Figure 4B). These observations were corroborated by immunofluorescence staining in A375 cells and confocal microscopy analysis (Figure 4C). Moreover, immunoblotting of histones isolated from a USP7 knockout (*USP7*-KO) A375 cell line that we generated using CRISPER-Cas9 technology also detected a decrease of the level of EZH2 and H3K27me3 and an increase level of H2BK120ub1 (Figure 4D), further supporting a notion that USP7 and PRC2 are functionally connected and that USP7 is a prerequisite for H3K27me3.

**Figure 4.**
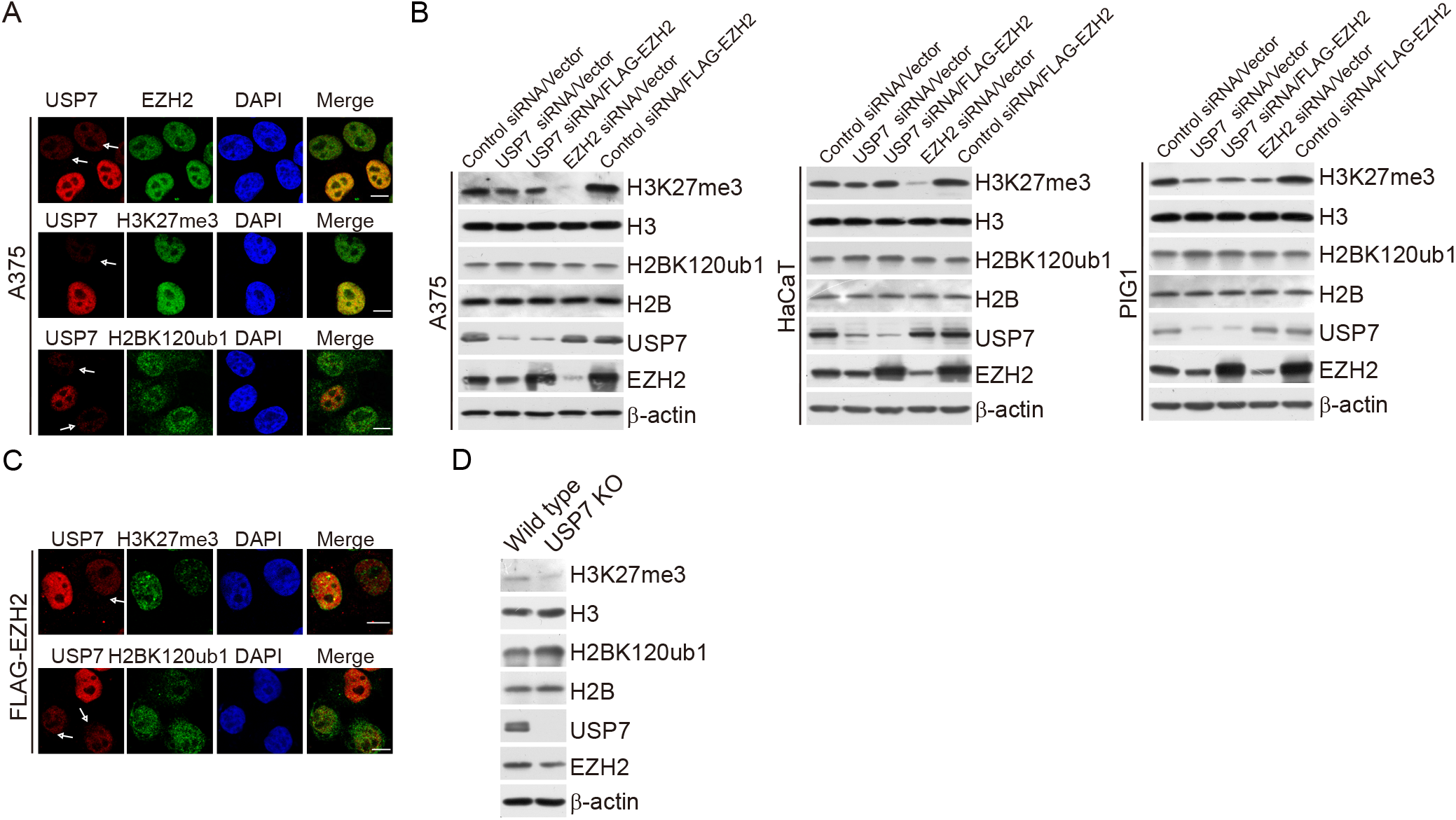
USP7 deubiquitinating H2BK120ub1 regulates the trimethylation of H3K27. (A) A375 cells transfected with *USP7* siRNA were fixed and immunostained with the indicated antibodies followed by confocal microscopy analysis. The white arrows indicate cells with USP7 depletion to different extents. Scale bar, 10 μm. (B) A375, HaCat, and PIG1 cells co-transfected with control siRNA or *USP7* siRNAs and Vector or FLAG-EZH2. Cellular extracts were collected and analyzed by western blotting. (C) A375 cells stably expressing FLAG-EZH2 transfected with *USP7* siRNA, then fixed, and immunostained with the indicated antibodies followed by confocal microscopy analysis. The white arrows indicate cells with USP7 depletion to different extents. Scale bar, 10 μm. (D) Histone extracts and cellular lysates from wild-type (WT) and *USP7* knockout (KO) A375 cells analyzed by western blotting.

### Genome-wide Identification of the Transcriptional Targets of the USP7/PRC2 Complex

To explore the biological significance of the physical association of USP7 with PRC2 complex, we analyzed the genome-wide transcriptional targets of USP7-PRC2 by ChIP-seq in A375 cells. At a *P-*value cutoff of 10^-3^, 19,189 USP7-specific binding peaks and 18,771 EZH2-specific binding sites were called (Figure 5A). Analysis of the overlapping sequences identified 1,236 specific promoter-binding genes of USP7 (5,361 genes) and EZH2 (3,452 genes) (Figure 5B). We then classified these co-targets into cellular signaling pathways including the Wnt, Hippo, MAPK, FoxO, Ras, oxytocin, and p53 pathways that are associated with tumorigenesis (Figure 5B). Significantly, EZH2 was significantly enriched in regions surrounding USP7 genomic binding sites (Figure 5C), and the genomic signatures analysis demonstrated that USP7 and EZH2 harbor similar binding motifs (Figure 5D). Collectively, these results support that USP7 and EZH2 physically interact and are functionally linked. Next, we sought to confirm whether USP7 is required for the chromatin recruitment of PRC2 to influence H3K27me3. Consistent with our previous experimental results, ChIP-seq showed that USP7 depletion led to a moderate reduction of H3K27me3 on USP7-occupied genes and stably overexpressed EZH2 could slightly up-regulate H3K27me3 (Figure 5E).

**Figure 5.**
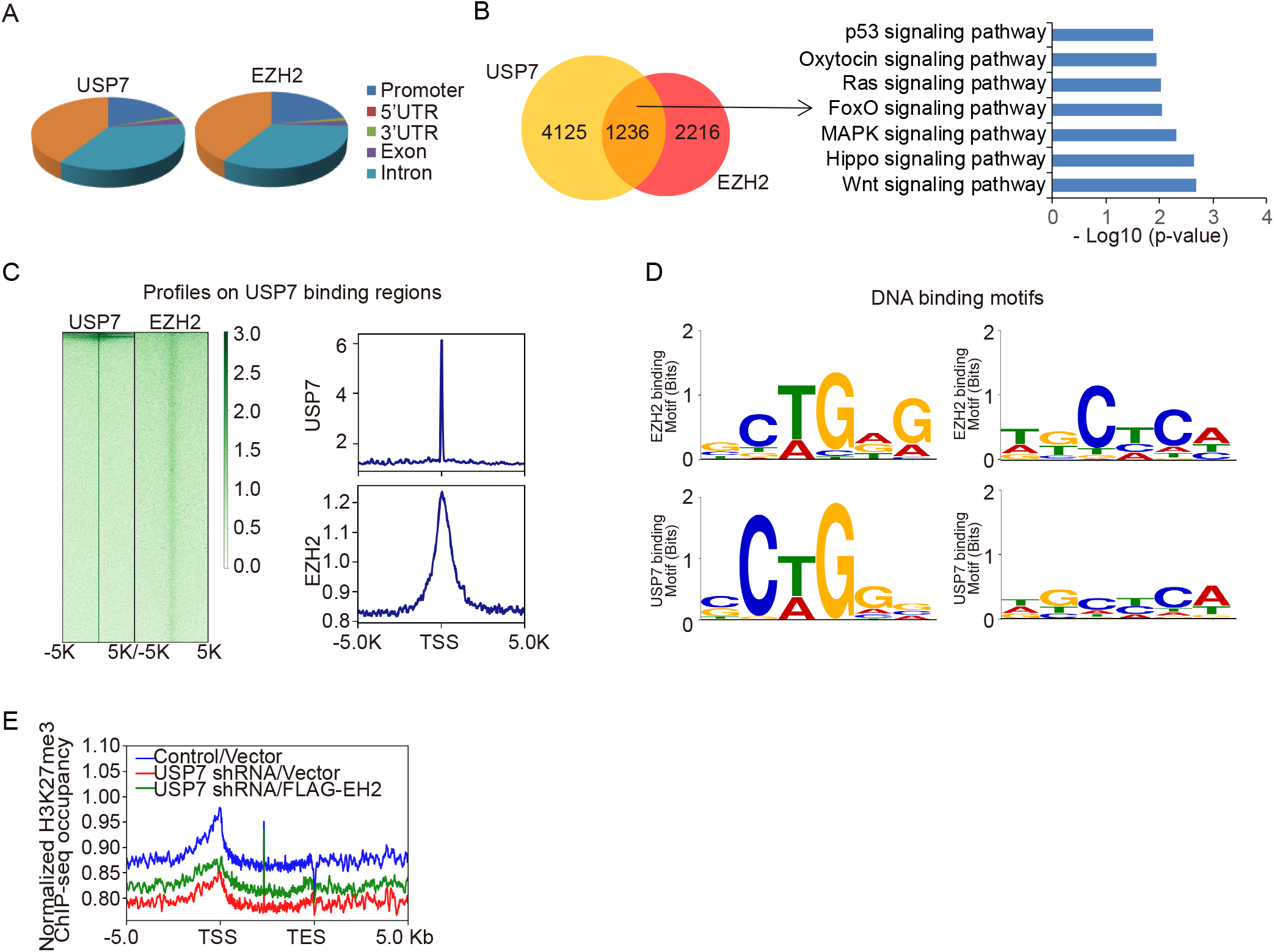
Genome-wide the transcription target analysis for USP7/PRC2 complex. (A) Genomic distribution of USP7 and EZH2 determined by ChIP-seq analysis. (B) Venn diagram of overlapping promoters bound by USP7 and EZH2 in A375 cells. The numbers represent the number of promoters targeted by the indicated proteins. The clustering of the 1,236 overlapping target genes of USP7/EZH2 into functional groups is shown. Detailed results of the ChIP-seq experiments are summarized in Supplementary Table 2. (C) ChIP-seq density heatmaps and profiles of EZH2 on USP7-binding sites. (D) MEME analysis of the DNA-binding motifs of USP7 and EZH2. (E) Average occupancy of H3K27me3 of the USP7/EZH2-occupied genes in A375 cells stably expressing Control shRNA, USP7 shRNA, or USP7 shRNA/FLAG-EZH2.

To further validate the ChIP-seq results, quantitative ChIP (qChIP) experiments were performed on selected genes, including *FOXO1, DUSP10, p21, CD82, BAI1, MOB2, DAB2IP, DLG3, HRK,* and *NISCH,* which showed that USP7 and EZH2 could indeed co-occupy the promoters of these genes, as well as H3K27me3 and H2BK120ub1 (Figure 6A). To further support the proposition that USP7 and PRC2 function as a protein complex at the target promoter, sequential ChIP or ChIP/Re-ChIP experiments were performed on the representative target genes *FOXO1* and *DUSP10*. The *FOXO1* and *DUSP10* promoters that immunoprecipitated with the antibodies against USP7 could be re-immunoprecipitated with antibodies against EZH2, SUZ12, EED, and H3K27me3 (Figure 6B).

**Figure 6.**
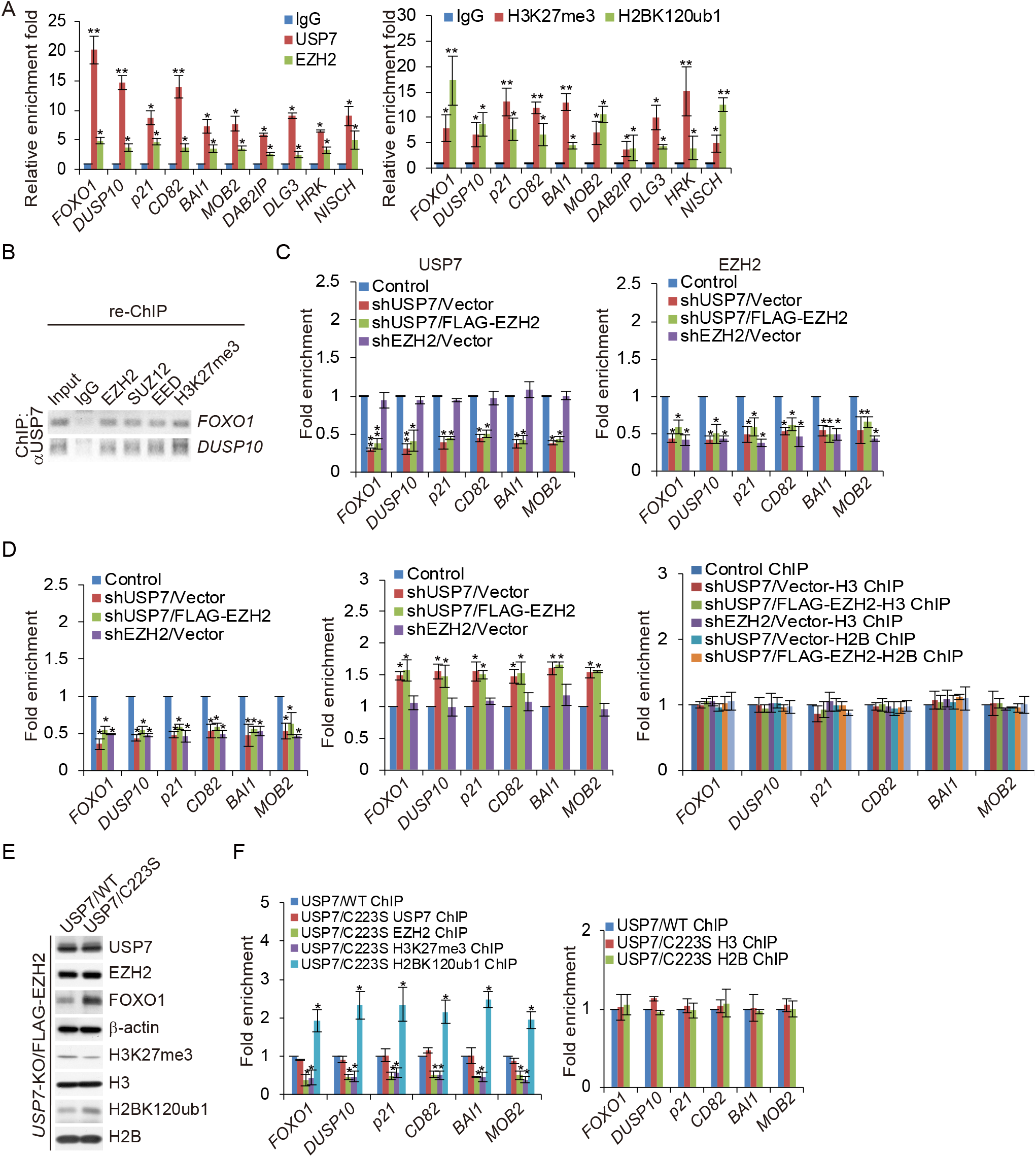
USP7 and EZH2 co-occupy target genes. (A) qChIP verification of the ChIP-seq results of the indicated genes with antibodies against the indicated proteins in A375 cells. (B) ChIP/Re-ChIP experiments on the promoters of the indicated genes with antibodies against the indicated proteins in A375 cells. (C-D) qChIP analysis of selected promoters in the A375 cells after co-transfection with indicated shRNA and Vector or FLAG-EZH2 using the indicated antibodies. Results are represented as the fold change over the control. (E) *USP7* KO A375 cells co-transfected with FLAG-EZH2 and USP7/WT, or catalytically inactive mutant of USP7 (USP7/C223S). Cellular and histone extracts were prepared and analyzed by western blotting. (F) USP7 knockout A375 cells co-transfected with FLAG-EZH2 and USP7/WT, or catalytically inactive mutant of USP7 (USP7/C223S). qChIP analysis of selected promoters was preformed using the indicated antibodies. Results are represented as fold change over USP7/WT. In A and C–F, each bar represents the mean ± SD from biological triplicate experiments. \**P* < 0.05 and \*\**P* < 0.01, by two-tailed unpaired Student’s t-test.

Since USP7 could promote the stability of EZH2 and USP7 depletion resulted in reduction of the H3K27me3 independent of protein level of EZH2, we next investigated the effect on USP7 depletion on the recruitment of USP7, EZH2, H3K27me3, and H2BK120ub1 to their co-target promoters. The qChIP analysis revealed that USP7 depletion reduced recruitment of USP7 to the promoters of these co-targets, which was not rescued by overexpression of EZH2, whereas EZH2 depletion had no effect on the recruitment of USP7 to its target promoters (Figure 6C). Consistently, the recruitment of EZH2 to its target promoters were both reduced in both USP7- or EZH2-depleted cells and co-expression of EZH2 with USP7 knockdown cells (Figure 6C). Consistent with our earlier observation, the level of H3K27me3 was obviously decreased at the candidate targets in USP7 and EZH2-depleted cells, which could not be recovered by stable overexpression of EZH2 in USP7-depleted cells (Figure 6D). In addition, the elevation of H2BK120ub1 enrichment on target promoters in USP7-deficient cells was not dependent on EZH2 (Figure 6D).

Ubiquitination of histone H2B was reported to directly regulate H3K4 methylation to activate gene transcription (42, 43), and USP7 forms a protein complex with guanosine 5’-monophosphate synthetase to catalyze the removal of H2BK120 ubiquitination (31). Thus, to explore whether USP7 deubiquitinated H2BK120ub for H3K27me3 independent of the protein level of EZH2, we co-transfected *USP7* knockout A375 cells with FLAG-EZH2 and USP7/WT or USP7/C223S. Western blotting analysis revealed that in *USP7* KO and EZH2 overexpression cells, USP7/WT, but not the catalytically inactive mutant of USP7 (USP7/C223S), could deubiquitinate H2BK120ub1 and restore the H3K27me3 level (Figure 6E). ChIP experiments further revealed that compared to USP7/WT cells, the recruitment of EZH2 and H3K27me3 to their co-target promoters was greatly reduced and the recruitment of H2BK120ub1 was elevated in USP7/C223S cells (Figure 6F). Overall, these results demonstrated that USP7 deubiquitinated H2BK120ub1, followed by recruiting the PRC2 complex to target promoters, supporting that USP7-regulated H2BK120ub1 deubiquitination and the physical interaction between USP7 and EZH2 are important for H3K27me3.

### Transcriptional Repression of Co-targets by the USP7/PRC2 Complex

In order to further explore the functional interaction between USP7 and PRC2 complex to regulate transcriptional repression of co-targets, qRT-PCR analysis and western blotting revealed that USP7 or EZH2 knockdown in A375 cells significantly increased the mRNA levels of *FOXO1, DUSP10, CD82,* and *p21* (Figure 7A and 7B), as well as the protein level of FOXO1, as a representative target (Figure 7A and 7B). Moreover, the increase of FOXO1 in USP7-depleted A375 cells could not be completely rescued by overexpression of EZH2, whereas EZH2 overexpression in USP7-proficient cells (transfected with control siRNA) significantly reduced the expression of FOXO1 (Figure 7C). In addition, IF analysis revealed that USP7 depletion could increase the level of FOXO1 even in cells overexpressing EHZ2 (Figure 7D), suggesting that USP7-catalyzed H2BK120ub1 deubiquitination is important for PRC2-mediated H3K27me3.

**Figure 7.**
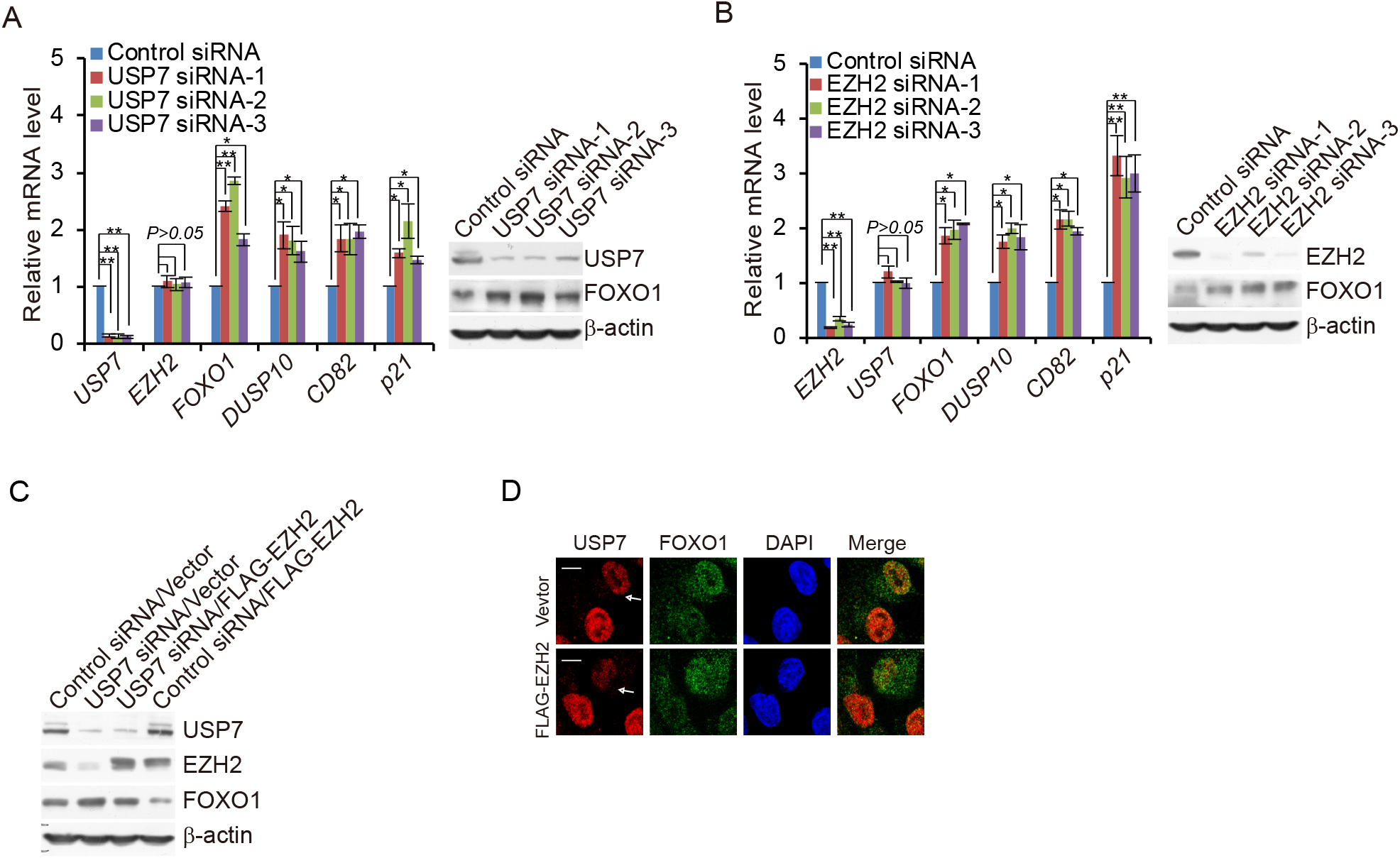
USP7 associated with EZH2 to co-suppress the transcription of targets. (A) A375 cells transfected with control siRNA or different sets of USP7 siRNAs were collected and analyzed by quantitative reverse transcription (qRT)-PCR and western blotting, respectively. (B) A375 cells transfected with control siRNA or different sets of *EZH2* siRNAs were collected and analyzed by qRT-PCR and western blotting, respectively. (C) A375 cells were co-transfected with control siRNA or *USP7* siRNAs and Vector or FLAG-EZH2; cellular extracts were collected and analyzed by western blotting. (D) A375 cells stably expressing FLAG-EZH2 were transfected with *USP7* siRNA, then fixed and immunostained with the indicated antibodies, followed by confocal microscopy analysis. The white arrows indicate cells with USP7 depletion to different extents. Scale bar, 10 μm.

### The USP7/EZH2-FOXO1 Axis Is Implicated in Melanoma Cell Proliferation and Melanomagenesis

To further explore the biological function of the USP7/PRC2 complex in melanoma progression, we respectively generated five types of A375 cell lines with control lentivirus and a lentivirus carrying *USP7* shRNA, *EZH2* shRNA, *USP7* shRNA and *FOXO1* shRNA, or *EZH2* shRNA and *FOXO1* shRNA. The efficiency of knockdown in these cells was verified by western blotting (Figure 8A). In addition, the EdU cell proliferation assay revealed that either knockdown of USP7 or EZH2 could inhibit cell proliferation, which could be recovered by concomitant knockdown of FOXO1 (Figure 8B). Moreover, growth curve measurement indicated that the growth delay induced by USP7 or EZH2 deficiency could be ameliorated by FOXO1 knockdown (Figure 8C). Colony formation assays showed that USP7 or EZH2 depletion hampered the colony formation of melanoma A375 cells, which was also rescued by knockdown of FOXO1 (Figure 8D). Treatment of these cells with different doses of the chemotherapy drugs (BRAF inhibitor) dabrafenib or vemurafenib used in melanoma treatment revealed that USP7- or EZH2-deficient cells were more sensitive to chemotherapeutics, which could be rescued by FOXO1 knockdown (Figure 8D).

**Figure 8.**
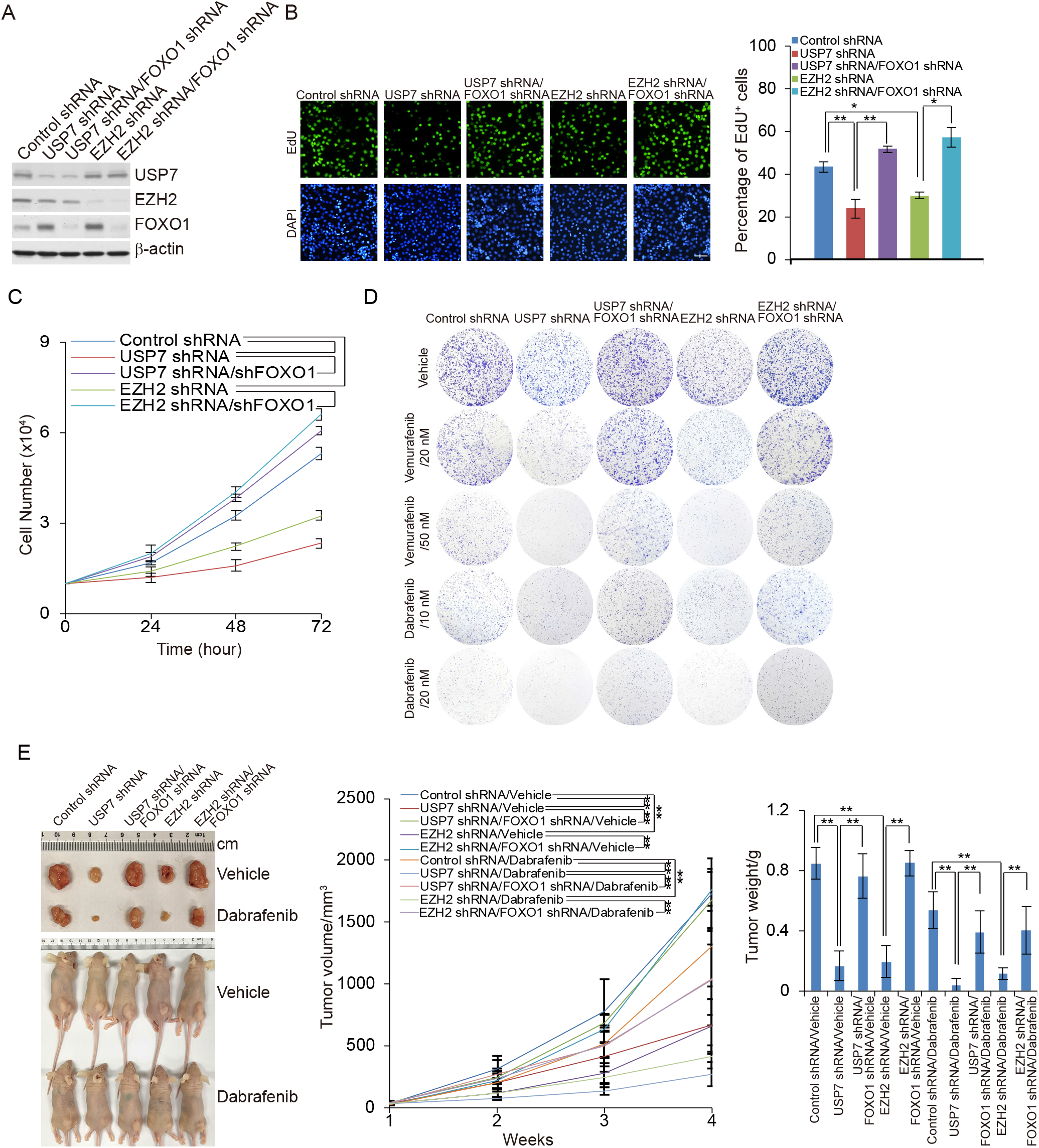
The USP7/EZH2-FOXO1 signaling pathway is required for melanoma cell proliferation and melanomagenesis. (A) A375 cells stably expressing the indicated shRNAs; the efficiency of knockdown was verified by western blotting. (B) EdU assays performed in A375 cells stably expressing the indicated shRNAs. Representative images and statistical analysis are shown. Scale bar, 50 μm. Each bar represents the mean ± SD from biological triplicate experiments. \**P* < 0.05, one-way ANOVA. (C) Growth curve assays in A375 cells stably expressing the indicated shRNAs. Each bar represents the mean ± SD from biological triplicate experiments. ** *P* < 0.01, two-way ANOVA. (D) A375 cells stably expressing the indicated shRNAs treated with different doses of dabrafenib or vemurafenib. Colony formation assays were performed, and the representative images are shown. (E) A375 tumors stably expressing indicated shRNAs were transplanted into athymic mice (n = 10); half of the mice in each group were randomly subjected to feed vehicle or dabrafenib daily one week post tumor injection. Representative tumors and sacrificed mice are shown (left panel). Tumor volumes were measured weekly and tumors were harvested and weighed when the mice were sacrificed. Each bar represents the mean ± SD. \**P* < 0.05, \*\**P* < 0.01, two-way ANOVA for tumor volume analysis and one-way ANOVA for tumor weight analysis. (right panel).

To confirm these results *in vivo*, tumors developed from these five A375 cell lines were ectopically transplanted into 6–8-week-old athymic mice (BABL/c nude; Charles River Laboratories). Tumor growth was substantially decreased in USP7-deficient or EZH2-deficient tumors, which could be alleviated by FOXO1 knockdown (Figure 8E). Moreover, FOXO1 depletion alleviated the increased sensitivity of USP7-deficient or EZH2-deficient tumors to dabrafenib (Figure 8E). Thus, USP7 recruiting PRC2 complex to suppress the expression of FOXO1 can protect melanin tumors from chemical toxicity, thus promoting the progression of melanoma.

### USP7 Is Implicated in Melanomagenesis and Poor Survival of Cancer Patients

FOXO1 as a tumor suppressor has been associated with multiple types of malignancies, including colon, cervical, prostate, gastric, and breast cancers (40, 44-47). Based on our findings that USP7 deubiquitinated H2BK120ub1 and recruited the PRC2 complex to suppress the transcription of FOXO1, we postulated that the USP7/EZH2-FOXO1 signaling pathway plays an important role in carcinogenesis. To validate this hypothesis, we first analyzed the protein levels of USP7, EZH2, and FOXO1 in human tissue arrays, including a series of paired tumor and adjacent normal tissue samples from the breast, kidney, lung, lymph node, ovary, esophagus, and skin. Immunohistochemical staining (IHC) analysis demonstrated the clear upregulation of USP7 and EZH2 accompanied by the downregulation of FOXO1 in these cancer tissues (Figure 9A). Next, we analyzed the expression profiles of USP7, EZH2, and FOXO1 in human tissue arrays of paired malignant and normal samples from patients with nevus, malignant melanoma, and metastatic melanoma. The IHC staining and the quantitation analysis revealed that compared to the normal skin tissue, USP7 and EZH2 were highly expressed but FOXO1 was expressed at a lower level in melanoma samples, and the expression levels of USP7 with EZH2 or FOXO1 were correlated with the histological grades of the tumor (Figure 9B). Moreover, the integrated cancer microarray database Oncomine (48) indicated that *USP7* mRNA levels are strikingly increased in benign melanocytic skin nevus and melanoma samples (Figure 9C). Importantly, Kaplan-Meier survival analysis from The Cancer Genome Atlas (TCGA) datasets showed that either a high expression level or increased copy number of *USP7* correlated with poor survival in melanoma patients (Figure 9D and 9E). Together, these findings are consistent with a role for USP7 in promoting melanoma.

**Figure 9.**
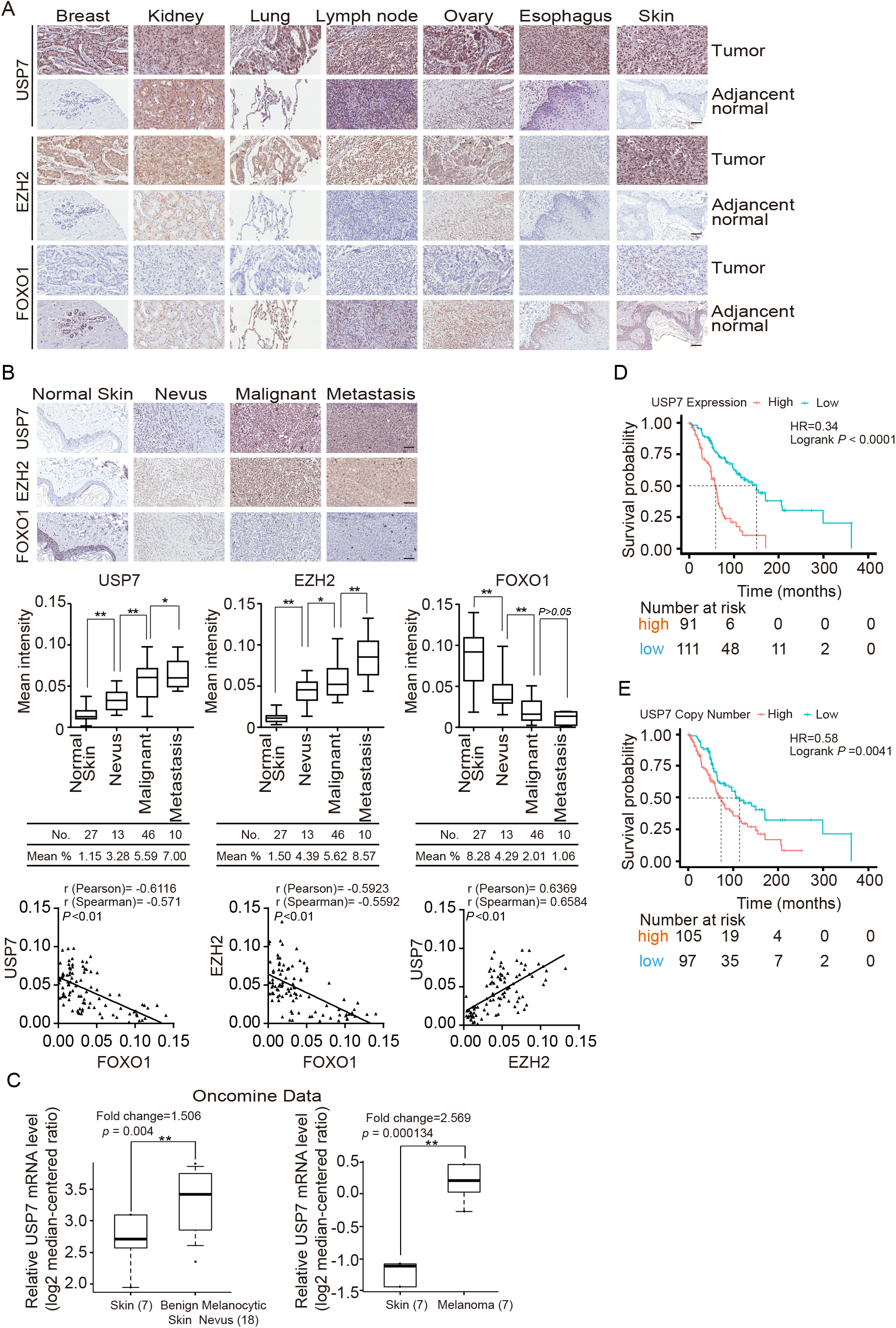
USP7 is implicated in melanomagenesis and poor patient survival. (A) Expression profiles of USP7, EZH2, and FOXO1 in human tissue arrays, including series of paired normal and tumor samples. Representative images (200× magnification) from three paired samples in each case are shown. Scale bar, 50 μm. (B) Human tissues containing benign nevus, malignant melanoma, metastatic melanoma, and tumor adjacent cervical samples were analyzed by immunohistochemical staining. Representative images (200× magnification) are shown. Scale bar, 50 μm. The staining intensity was determined by Image-Pro Plus software and presented as box plots. \**P* < 0.05; \*\**P* < 0.01; one-way ANOVA. The correlation coefficient and *P* values were analyzed. (C) Analysis of Talantov melanoma (left panel) or Haqq melanoma (right panel) from Oncomine for the expression of *USP7* in normal human skin tissues and benign melanocytic skin nevus samples or in normal human skin tissues and melanoma samples. Data are presented as box plots (\*\**P* < 0.01). (D) Kaplan-Meier survival analysis for the relationship between survival time of melanoma patients and the mRNA expression level of *USP7* with survival packages from https://tcga-data.nci.nih.gov/docs/publications/tcga/. (E) Kaplan-Meier survival analysis of the relationship between survival time of melanoma patients and gene copy number status of *USP7* with survival packages from https://tcga-data.nci.nih.gov/docs/publications/tcga/.

## DISCUSSION

Dynamic regulation of epigenetic modifiers plays a key role in the modulation of gene expression and consequently fate specification (49, 50). In particular, increased activity of the histone methyltransferase EZH2 has been associated with various cancers, including melanoma (8). In this study, we showed that EZH2 is physically associated with and is stabilized by the deubiquitinase USP7. Importantly, USP7 functions as an alternate histone modification enzyme responsible for H2BK120ub1 deubiquitination (31), recruiting PRC2 complex to the chromatin, providing a molecular basis for the interplay of H2BK120ub and H3K27me3 in chromatin remodeling. To date, PRC2 has been shown to mediate transcriptional repression by distinct sequence-specific transcription factors (51). Interestingly, we found that FOXK1 could also physically interact with USP7 and the PRC2 complex in A375 melanoma cells, suggesting that USP7 and PRC2 may share similar transcription factors and thus respond to the same signal pathways. However, further investigations are needed to explore the scope and variety of the functionality of the USP7/PRC2 complex and to determine whether this functionality involves additional elements.

The elevated expression of EZH2 often correlates with a poor prognosis in cancer patients (52, 53). However, the mechanism of EZH2 overexpression in cancers has remained elusive. Indeed, the regulators of EZH2 expression are also critical factors for tumorigenesis. For example, Myc binds to the *EZH2* promoter and directly activates its transcription, and ANCCA, a co-activator of androgen receptor, can enhance E2F-mediated EZH2 transcription (54). Accumulating evidence indicates that the activity and stability of EZH2 are also regulated by post-translational modifications such as through CDK1/2, which could phosphorylate EZH2 at multiple sites (55), and JAK2, which phosphorylates EZH2 at tyrosine 641 to promote the interaction of EZH2 with β-TrCP and the consequent degradation of EZH2 (56). However, the protein that guarantees the stability of EZH2 has not been identified until now. USP7 has been reported to stabilize a number of proteins, thereby playing a role in multiple cellular processes (24, 25). Given our finding that USP7 is physically associated with the PRC2 complex, it is reasonable that USP7 would stabilize EZH2 through its deubiquitinase activity. Future investigations will determine whether and how the USP7/PRC2 complex functions in normal development, as well as in the development and progression of tumors originating from other tissues.

Interestingly, like many other USP members, USP7 has been reported to deubiquitinate H2BK120ub1. Recent studies have shown that H2BK120ub1 is a prerequisite for H3K4me3 (57). Moreover, in embryonic stem cells, bivalent chromatin domains containing H3K4me3 and H3K27me3 marks silence developmental genes, while keeping them poised for activation following differentiation (58). Recently, some cancer types have been shown to exhibit partial recapitulation of bivalent chromatin modifications (59), but the roles of deubiquitinated histone H2B in histone cross-talk are poorly understood. Our findings suggest that USP7 provides a molecular basis for the interplay of H2BK120ub1 and H3K27me3 in chromatin remodeling, which is independent of the protein level of EZH2. However, the ChIP-seq results clearly showed incomplete overlap between USP7 and EZH2 targets, suggesting that the PRC2 complex could recruit some other transcriptional cofactors. USP7 could also deubiquitinate non-histone proteins with independent functions under certain physiological states.

The FOXO1 transcription factor orchestrates the regulation of genes involved in the apoptotic response (60), cell cycle checkpoints (61), and cellular metabolism (62). FOXO1 is a putative tumor suppressor, and its gene expression is dysregulated in some cancers, including endometrial cancer (38) and melanoma (63). Consistent with our observation that USP7 coordinated with the PRC2 complex is linked to downregulation of FOXO1, we showed that the expression level of FOXO1 is decreased and negatively associated with those of USP7 and EZH2 in various tumor tissue samples. Moreover, we demonstrated that FOXO1 loss of function could rescue EZH2 or USP7 depletion-induced phenotypes *in vitro* and *in vivo*. Together, these findings provide a molecular basis for understanding the dysregulation of FOXO1 in melanoma.

Acquired resistance is a major problem limiting the long-term effectiveness of targeted cancer therapeutics (64, 65). Resistance can be acquired through secondary mutations or amplification of the primary drug targets, or by activation of bypass signaling pathways (66–68). High EZH2 expression was shown to be associated with more malignant forms of melanoma; thus, patients with EZH2 high expression showed significantly shorter overall survival as compared with those with lower EZH2 expression. Furthermore, a higher non-synonymous EZH2 mutation frequency was identified in melanoma compared to that in other solid tumors. In particular, cutaneous melanoma was the only solid cancer with non-synonymous mutations affecting tyrosine 646* (Y646*) in the SET domain of EZH2, which has been implicated in the haematopoietic system (8). Thus, an EZH2 inhibitor is not a suitable alternative for melanoma treatment. Since both USP7 and EZH2 were significantly overexpressed in a subset of patients with melanoma, and the levels of these two factors were positively correlated, our results suggest that targeting USP7 may provide an effective treatment for patients with melanoma under a biotherapy strategy, particularly in light of recent efforts to develop a specific small-molecule antagonist toward the catalytic activity of USP7. Moreover, as USP7 not only has an effect on the EZH2 protein level but also plays a key role in the function of EZH2, it could be a promising therapeutic target to replace EZH2, at least for the treatment of melanoma with BRAF inhibitor resistance.

In summary, we have demonstrated that USP7 is physically associated with the PRC2 complex, promoting EZH2 stabilization, and chromatin engagement of the PRC2 complex to ultimately drive the development of melanoma. Our findings support a model by which upregulation of USP7 in melanoma results in an elevated abundance of EZH2, as well as H2BK120ub1 removal and an increased level of H3K27me3. Although it’s hard to specify whether USP7 is in dependent of or in concert with stabilizing EZH2 in PRC2 recruitment, our data indicates that USP7 coordinates with the PRC2 complex to promote tumorigenesis, supporting the pursuit of USP7 as a potential target for melanoma intervention.

## METHODS

### Antibodies and reagents

The sources of antibodies against the following proteins were as follows: HA (sc-805) from Santa Cruz Biotechnology; β-actin (A1978), EZH2 (AV38470, for IHC), and FLAG (F3165) from Sigma; USP7 (05-1946, for WB, IF, and IP) from Millipore; Ubiquityl-Histone H2B (K120) (#5546, for WB, IF, and ChIP) and FOXO1 (#2880, for WB and IHC) from Cell Signaling Technology; SUZ12 (ab175187, for WB and IP), H3K27me3 (ab6002, for WB, IF, ChIP, and ChIP-seq), H2B (ab64165, for WB and ChIP), and histone H3 (ab1791, for WB and ChIP) from Abcam; USP7 (A300-033A, IP, IF, IHC, and ChIP) from Bethyl Lab; Myc (M047-3) from MBL; EED (GTX33168, for WB and IP) from GeneTex; EZH2 (612667, for WB, IP, and IF) from BD Transduction Laboratories; and EZH2 (39901, for ChIP and ChIP-seq) from Active Motif. Anti-HA affinity gel (E6779), anti-FLAG M2 affinity gel (A2220), 3× FLAG peptide (F4799), MG132 (SML1135), blasticidin (15205), puromycin (P8833), and doxycycline (D9891) were purchased from Sigma. Ni-NTA Purification System (K950-01) was purchased from Invitrogen. CHX and HBX 41,108 were purchased from TOCRIS. GNE-6640 was purchased from Glixx Laboratories. Dabrafenib (GSK2118436) and venurafenib were purchased from Selleck.

### Plasmids

The FLAG- or Myc-tagged USP7/WT was amplified from *USP7* cDNA kindly provided by Dr. Yang Shi (Harvard Medical School, Boston, MA, USA) and Dr. Ruaidhri J. Carmody (University of Glasgow, Scotland, UK) and integrated into pLVX-Tight-Puro, pLenti-hygro, or pcDNA3.1 vector, while the FLAG-tagged USP7/C223S carried by pLVX-Tight-Puro or pLenti-hygro vector was generated by a quick-change point mutation assay. His-tagged USP7/WT, USP7/C223S, and deletion mutants of USP7 were carried by pFastBac-HTA vector. CRISPR/Cas9 constructs lentiCas9-Blast (Addgene plasmid # 52962) and lentiGuide-Puro (Addgene plasmid # 52963) were gifts from Dr. Feng Zhang (Broad Institute, Cambridge), and HA-tagged ubiquitin K48-only (Plasmid #17605) and K63-only (Plasmid #17606) were gifts from Dr. Ted Dawson (Johns Hopkins University School of Medicine, Baltimore, MD, USA). FLAG-tagged EZH2 carried by pLenti-hygro or pcDNA3.1 vector, GST-tagged deletion mutants of EZH2 in pGEX4T-3 vector, and FLAG-tagged SUZ12 or EED in pcDNA3.1 vector were purchased from Youbio (Hunan, China).

### Cell culture

HEK293T, A375, PIG1, and Sf9 cells were obtained from American Type Culture Collection (Manassas, VA, USA) and cultured according to the manufacturer’s instructions. HaCAT cells were from the China Center for Type Culture Collection (CCTCC) and cultured according to the manufacturer’s instructions. Cell lines with Dox-induced protein expression were established in two steps. First, the cells were infected with a lentivirus carrying rtTA and subjected to neomycin selection. Subsequently, the established rtTA cells were infected with a virus carrying the pLenti-Tight-Puro vector encoding USP7/wt or USP7/C223S followed by puromycin selection. All of the cells integrated with rtTA were cultured in Tet Approved FBS and medium from Clontech. All the cells were authenticated by examination of morphology and growth characteristics and confirmed to be mycoplasma-free.

### *USP7* KO cell generation

USP7 KO A375 cells were generated by co-transfection of the plasmid encoding FLAG-Cas9 (lentiCas9-Blast) and sgRNA plasmid (lentiGuide-Puro) targeting *USP7* (AATCAGATTCAGCATTGCAC). Forty-eight hours after transfection, the cells were selected by blasticidin (5 μg/ml) and puromycin (1 μg/ml) for 2 days.

### Immunopurification and silver staining

Lysates from A375 cells stably expressing FLAG-EZH2 with or without Dox-inducible expression of FLAG-USP7 were prepared by incubating the cells in lysis buffer containing protease inhibitor cocktail (Roche). Anti-FLAG immunoaffinity columns were prepared using anti-FLAG M2 affinity gel (Sigma) following the manufacturer’s protocols. Cell lysates were obtained from approximately 5 × 10^7^ cells and applied to an equilibrated FLAG column of a 1-ml bed volume to allow for adsorption of the protein complex to the column resin. After binding, the column was washed with cold PBS plus 0.2% Nonidet P-40. FLAG peptide (Sigma) was applied to the column to elute the FLAG protein complex according to the manufacturer’s instructions. The elutes were collected and visualized on a NuPAGE 4–12% Bis-Tris gel (Invitrogen) followed by silver staining with silver staining kit (Pierce). The distinct protein bands were retrieved and analyzed by LC-MS/MS.

### Nano-HPLC-MS/MS analysis

LC-MS/MS analysis was performed using a Thermo Finnigan LTQ linear ion trap mass spectrometer in line with a Thermo Finnigan Surveyor MS Pump Plus HPLC system. Generated tryptic peptides were loaded onto a trap column (300SB-C18, 5 × 0.3 mm, 5-μm particle size; Agilent Technologies, Santa Clara, CA, USA) connected through a zero-dead-volume union to the self-packed analytical column (C18, 100 μm i.d × 100 mm, 3-μm particle size; SunChrom, Germany). The peptides were then eluted over a gradient (0–45% B in 55 min, 45–100% B in 10 min, where B = 80% acetonitrile, 0.1% formic acid) at a flow rate of 500 nl/min and introduced online into the linear ion trap mass spectrometer (Thermo Fisher Corporation, San Jose, CA, USA) using nano electrospray ionization. Data-dependent scanning was incorporated to select the five most abundant ions (one microscan per spectrum; precursor isolation width, 1.0 m/z; 35% collision energy, 30 ms ion activation; exclusion duration, 90 s; repeat count, 1) from a full-scan mass spectrum for fragmentation by collision-induced dissociation. MS data were analyzed using SEQUEST (v. 28) against the NCBI human protein database (downloaded December 14, 2011; 33,256 entries), and results were filtered, sorted, and displayed using the Bioworks 3.2. Peptides (individual spectra) with Preliminary Score (Sp) ≥ 500; Rank of Sp (RSp) ≤ 5; and peptides with +1, +2, or +3 charge states were accepted if they were fully enzymatic and had a cross correlation (Xcorr) of 1.90, >2.75, and >3.50, respectively. At least two distinct peptides were assigned to each identified protein. The following residue modifications were allowed in the search: carbamidomethylation on cysteine as a fixed modification and oxidation on methionine as variable modification. Peptide sequences were searched using trypsin specificity and allowing for a maximum of two missed cleavages. Sequest was searched with a peptide tolerance of 3 Da and a fragment ion tolerance of 1.0 Da.

### FPLC chromatography

A375 nuclear extracts and FLAG-USP7-containing protein complexes were applied to a Superpose 6 size exclusion column (GE Healthcare) that had been equilibrated with dithiothreitol-containing buffer and calibrated with protein standards (Amersham Biosciences). The column was eluted at a flow rate of 0.5 ml/min and fractions were collected.

### Immunoprecipitation

Cell lysates were prepared by incubating the cells in NETN buffer (50 mM Tris-HCl, pH 8.0, 150 mM NaCl, 0.2% Nonidet P-40, 2 mM EDTA) in the presence of protease inhibitor cocktails (Roche) for 20 min at 4°C, followed by centrifugation at 14,000 ×*g* for 15 min at 4°C. For IP, approximately 500 μg of protein was incubated with control or specific antibodies (1–2 μg) for 12 h at 4°C with constant rotation; 50 μl of 50% protein G magnetic beads (Invitrogen) was then added and the incubation was continued for an additional 2 h. The beads were washed five times using the lysis buffer. Between washes, the beads were collected by a magnetic stand (Invitrogen) at 4°C. The precipitated proteins were eluted from the beads by re-suspending the beads in 2× SDS-PAGE loading buffer and boiling for 5 min. The boiled immune complexes were subjected to SDS-PAGE followed by IB with appropriate antibodies.

### In vivo deubiquitination assay

Cells with different treatments were lysed in RIPA buffer containing 50 mM Tris-HCl (pH 7.4), 150 mM NaCl, 1% NP-40, 0.1% SDS, and protease inhibitor at 4°C for 30 min with rotation, and centrifuged at 20,000 ×*g* for 15 min. Approximately 0.5–1.5 mg of cellular extracts were immunoprecipitated with anti-FLAG agarose affinity gel for 2 h. The beads were then washed five times with RIPA buffer, boiled in SDS loading buffer, and subjected to SDS-PAGE followed by IB.

### Recombinant protein purification and pull-down assays

Recombinant baculovirus carrying full-length USP7/wt or deletion mutants of *USP7* were generated with the Bac-to-Bac System (Invitrogen). Infected Sf9 cells were grown in spinner culture for 48–96 h at 27°C and His-tagged protein-purified using Ni^2+^-NTA agarose (Invitrogen) according to standard procedures. For the His pull-down assay, His-tagged protein was incubated with recombinant EZH2, SUZ12, or EED that was *in vitro-*transcribed and translated according to the manufacturer’s procedures (TNT T7 Quick Coupled Transcription/Translation Kit; Promega, Leiden, the Netherlands) at 4°C overnight. GST-fusion proteins were purified from *Escherichia coli* by glutathione-Sepharose 4B beads (GE Healthcare) and then washed with high salt buffer (20 mM Tris-HCl pH 7.4, 0.1 mM EDTA, and 300 mM NaCl). For the GST pull-down assay, GST-fusion proteins were incubated with *in vitro*-transcribed and translated proteins at 4°C overnight. The beads were washed three times, then boiled in SDS loading buffer, and subjected to SDS-PAGE followed by immunoblotting.

### RNA interference

All siRNA transfections were performed using Lipofectamine RNAi MAX (Invitrogen) following the manufacturer’s recommendations. The final concentration of the siRNA molecules was 10 nM, and the cells were harvested 72–96 h later according to the experimental purpose. The *USP7* siRNA was a mixture of individual siRNAs against *USP7*, and the *EZH2* siRNA was a mixture of individual siRNAs against *EZH2*. The siRNA sequences are shown in Supplementary Table 3.

### Lentiviral production

The shRNAs targeting *USP7* (Sigma), *EZH2*, or *FOXO1* in pLKO vector or vectors encoding rtTA, USP7, EZH2 carried by pLenti vectors, as well as three assistant vectors (pMDLg/pRRE, pRSV-REV, and pVSVG) were transiently transfected into HEK293T cells. Viral supernatants were collected 48 hours later, clarified by filtration, and concentrated by ultracentrifugation. The shRNA sequences are shown in Supplementary Table 4.

### qRT-PCR

Total cellular RNA was isolated with TRIzol reagent (Invitrogen) and used for first-strand cDNA synthesis with the Reverse Transcription System (Roche). Quantitation of all gene transcripts was performed by qPCR using a Power SYBR Green PCR Master Mix (Roche) and an ABI PRISM 7500 sequence detection system (Applied Biosystems) with the expression of *GAPDH* as the internal control. The qRT-PCR primers are shown in Supplementary Table 5.

### Chromatin Immunoprecipitation

Approximately 10 million cells were cross-linked with 1% formaldehyde for 10 min at room temperature and quenched by the addition of glycine to a final concentration of 125 mM for 5 min. The fixed cells were resuspended in SDS lysis buffer (1% SDS, 5 mM EDTA, 50 mM Tris-HCl pH 8.1) in the presence of protease inhibitors and subjected to 3 × 10 cycles (30 seconds on and 30 seconds off) of sonication (Bioruptor, Diagenode) to generate chromatin fragments of ∼300 bp in length. Lysates were diluted in buffer containing 1% Triton X-100, 2 mM EDTA, 20 mM Tris-HCl pH 8.1, 150 mM NaCl. For IP, the diluted chromatin was incubated with control or specific antibodies (2 μg) for 12 hours at 4°C with constant rotation; 50 μl of 50% protein G magnetic beads was then added and the incubation was continued for an additional 2 hours. The beads were washed with the following buffers: TSE I (0.1% SDS, 1% Triton X-100, 2 mM EDTA, 20 mM Tris-HCl pH 8.1, 150 mM NaCl), TSE II (0.1% SDS, 1% Triton X-100, 2 mM EDTA, 20 mM Tris-HCl pH 8.1, 500 mM NaCl), buffer III (0.25 M LiCl, 1% NP-40, 1% sodium deoxycholate, 1 mM EDTA, 10 mM Tris-HCl pH 8.1), and Tris-EDTA buffer. Between washes, the beads were collected by a magnetic stand at 4°C. The pulled-down chromatin complex together with the input were de-crosslinked at 70°C for 2 hours in elution buffer (1% SDS, 5 mM EDTA, 20 mM Tris-HCl pH 8.1, 50 mM NaCl, 0.1 mg/ml Proteinase K). Eluted DNA was purified with a PCR purification kit (Qiagen) and analyzed by qPCR.

### ChIP sequencing, qChIP and ChIP/Re-ChIP

A375 cells or A375 cells stably expressing Control shRNA, USP7 shRNA, and USP7 shRNA/FLAG-EZH2 were maintained in DMEM supplemented with 10% FBS. Approximately 5 × 10^7^ cells were used for each ChIP-Seq assay. The chromatin DNA was precipitated by polyclonal antibodies against IgG, USP7, EZH2, or H3K27me3. The DNA was purified with a Qiagen PCR purification kit. In-depth whole-genome DNA sequencing was performed by BGI, Beijing. The raw sequencing image data were examined using the Illumina analysis pipeline, aligned to the unmasked human reference genome (UCSC GRCh37, hg19) using Bowtie2, and further analyzed by MACS2 (Model-based Analysis for ChIP-Seq, https://github.com/taoliu/MACS). Enriched binding peaks were generated after filtering through the input data. The genomic distribution of USP7- and EZH2-binding sites was analyzed by ChIPseeker, an R package for ChIP peak annotation, comparison, and visualization (http://bioconductor.org/packages/release/bioc/html/ChIPseeker.html). DNA-binding motifs of USP7 and EZH2 were analyzed by MEME (http://meme-suite.org/tools/meme). For visualization of ChIP-Seq data, we generated a bigwig track heatmap for genome browsers using deepTools (version 2.2.0). Eluted DNA was purified using a PCR purification kit (QIAGEN), and qChIPs were performed using the TransStart Top Green qPCR Supermix (TransGen Biotech) by qPCR on the ABI 7500-FAST System. Re-ChIPs were performed essentially in the same manner as the primary IPs. Bead eluates from the first immunoprecipitation were incubated with 10 mM DTT at 37°C for 30 min and diluted 1:50 in dilution buffer (1% Triton X-100, 2 mM EDTA, 150 mM NaCl, 20 mM Tris-HCl pH 8.1) followed by re-IP with the secondary antibodies. The final elution step was performed using 1% SDS solution in Tris-EDTA buffer, pH 8.0. The qChIP PCR primers are listed in Supplementary Table 6.

### Colony formation assay

A375 cells stably expressing the indicated shRNAs were treated with different doses of the chemotherapy drug dabrafenib or vemurafenib. The cells were then maintained for 14 days, fixed with methanol, and stained by crystal violet.

### Tissue specimens

The tissue samples were obtained from surgical specimens from patients with nevus, malignant melanoma, and metastasis melanoma. The samples were frozen in liquid nitrogen immediately after surgical removal and maintained at −80°C until protein extraction. Human skin tissues were prepared; incubated with antibodies against USP7, EZH2, or FOXO1; and processed for IHC with standard DAB staining protocols. Images for adjacent normal (27), nevus (13), malignant melanoma (46), and metastasis melanoma (10) samples were collected under microscopy with 200× magnification. The image quality was evaluated and the background with uneven illumination was corrected with Image-Pro Plus software. The normal skin cells, nevus cells, or melanoma cells were selected as the region of interest (ROI) according to the morphological features of the tissue or cells. The scores of the stained sections were determined by evaluating the mean intensity and nuclear staining extent of immunopositivity following the instructions of Image-Pro Plus software.

### Tumor xenografts

A375 cells were plated and infected *in vitro* with mock or lentiviruses carrying control shRNA, *USP7* shRNA, or *EZH2* shRNA together with *FOXO1* shRNA. Forty-eight hours after infection, 3 × 10^6^ viable A375 cells in 100 μl PBS were injected into the 6–8-week-old female athymic mice (BALB/c; Charles River, Beijing, China). One week after inoculation, the mice were subjected to a feed vehicle or dabrafenib (30 mg/kg body weight) daily for half the mice in each group, and then tumor growth and body weight were monitored over the following 3 weeks. Five animals per group were used in each experiment. Tumors were measured weekly using a Vernier caliper and the volume was calculated according to the formula: π/6 × length × width^2^. The measurement and data processing were performed in a blinded manner to the group.

### Statistics

Data from biological triplicate experiments are presented with error bar as mean ± SD. Two-tailed unpaired Student’s t-test was used for comparing two groups of data. Analysis of variance (ANOVA) with Bonferroni’s correction was used to compare multiple groups of data. A *P* value of less than 0.05 was considered significant. All the statistical testing results were determined by SPSS software. Before statistical analysis, variation within each group of data and the assumptions of the tests were checked.

### Study approval

All animal handling and experiments were approved by the Animal Care Committee of Capital Medical University. The collection and analysis of human tissue samples were approved by the Ethics Committee of the Capital Medical University, and informed consent was obtained from all patients.

## AUTHOR CONTRIBUTIONS

Dongxue Su and Lin Shan conceived this project; Dongxue Su, Wenjuan Wang, and Yongqiang Hou conducted experiments; Dongxue Su, Wenjuan Wang, Yongqiang Hou, Yue Wang, Chao Yang, Beibei Liu, Xing Chen, Xiaodi Wu, Jiajing Wu, Dong Yan, Shuqi Wei, and Lu Han acquired data; Dongxue Su, Liyong Wang, Wenjuan Wang, Yongqiang Hou, Shumeng Liu, Lei Shi, and Lin Shan analyzed data; Dongxue Su and Lin Shan wrote the manuscript.

## ACKNOWLEDGMENTS

This work was supported by grants (31871279 to L.S.) from the National Natural Science Foundation of China.

