## Supplemental Data 2 for "Bimodal Regulation of the PRC2 Complex by USP7 Underlies Melanomagenesis"

**Supplementary Table 3. siRNA sequences**

| siRNAs | Sequences |
| --- | --- |
| USP7-1 | GACGUUUCGAAUAGAGGAA |
| USP7-2 | GCACUAAUGCUUACAUGUU |
| USP7-3 | GACUUUGAGAACAGGCGAA |
| USP7 5’UTR-1 | CUCACCUCGUCAGCCACUA |
| USP7 5’UTR-2 | CAAGUCUUGUGUUUAGGCU |
| EZH2-1 | AGAATACTTGAACTTGTCC |
| EZH2-2 | GAGGTTCAGACGAGCTGAT |
| EZH2-3 | GCAACACCCAACACTTATAAG |

**Supplementary Table 4. Lentiviral shRNA sequences**

| shRNAs | Sequences |
| --- | --- |
| Control | CCGGGATATGGGCTGAATACAAACTCGAGTTTGTATTCAGCCCATATCTTTTTG |
| USP7 | CCGGCCTGGATTTGTGGTTACGTTACTCGAGTAACGTAACCACAAATCCAGGTTTTTG |
| EZH2 | CCGGGCAACACCCAACACTTATAAGCTCGAGCTTATAAGTGTTGGGTGTTGCTTTTTG |
| FOXO1 | CCGGCCAAACACCAGTTTGAATTCTCGAGAATTCAAACTGGTGTTTGGTTTTTG |

Note: Red color indicates the targeting sequence against the corresponding genes.

**Supplementary Table 5. qRT-PCR primers**

| **Genes** | **Forward Primer Sequences** | **Reverse Primer Sequences** |
| --- | --- | --- |
| *USP7* | ATTCCTAACATTGCCACCAG | ATTTACACCATTTGCCATCC |
| *EZH2* | GTACACGGGGATAGAGAATGTGG | GGTGGGCGGCTTTCTTTATCA |
| *FOXO1* | TCGTCATAATCTGTCCCTACACA | CGGCTTCGGCTCTTAGCAAA |
| *DUSP10* | ATCGGCTACGTCATCAACGTC | TCATCCGAGTGTGCTTCATCA |
| *p21* | CGATGGAACTTCGACTTTGTCA | GCACAAGGGTACAAGACAGTG |
| *CD82* | TGTCCTGCAAACCTCCTCCA | CCATGAGCATAGTGACTGCCC |
| *GAPDH* | GAAGGTGAAGGTCGGAGTC | GAAGATGGTGATGGGATTTC |

**Supplementary Table 6. qChIP primers**

| **Genes** | **Forward Primer Sequences** | **Reverse Primer Sequences** |
| --- | --- | --- |
| *FOXO1* | CAGCCCAGATGAGGAAAGG | CGGGAGATAGGACCAAAGC |
| *DUSP10* | TTCTTTAGGGTTCTTGGACG | AAAGTTGCCCTGGGATACA |
| *p21* | AGGTTCAAGCGATTCTCC | ATCACAGGGTCAGGAGTT |
| *CD82* | CCCTGACTTCTTTACAGTTTGC | GGAACTTGTTAGGAGTGCGTAT |
| *BAI1* | CACTGTGACGGGTAAATTATGG | ACCTCTGCCCTTCCTCTGC |
| *MOB2* | GTGTCTGTGCTGGGTTGGTG | ATGGGAGTTGTGGTGTCTGG |
| *DAB2IP* | GGGCAAATACTTGAAATGG | CCTCCCTGGTGACCTTGT |
| *DLG3* | GATTTGGGTTTGCGTGTT | GGAATGGAGGAAGGATGAC |
| *HRK* | ACCCTTACCTACCTGTGCCT | CCTTGCCTCTGCTGCCTA |
| *NISCH* | GAGCTACAAAGTCGTGGGC | TTATCTGGAAAGTGGAGGTCAT |
